# Crystal structure determination of the armadillo repeat domain of *Drosophila* SARM1 using MIRAS phasing

**DOI:** 10.1101/2021.01.31.428505

**Authors:** Weixi Gu, Zhenyao Luo, Clemens Vonrhein, Xinying Jia, Thomas Ve, Jeffrey D. Nanson, Boštjan Kobe

**Author notes:** These authors contributed equally to this work (Weixi Gu, Zhenyao Luo).

## Abstract

We describe the crystal structure determination of the ARM domain of *Drosophila* SARM1 (dSARM1^ARM^), which required combination of a number of sources of phase information in order to obtain interpretable electron density maps. SARM1 is a central executioner of the process of axon degeneration, a common feature of the early phase of a range of neurodegenerative diseases. SARM1 is held in the inactive state in healthy axons by its N-terminal auto-inhibitory ARM domain, and is activated to cleave NAD^+^ upon injury, triggering the subsequent axon degeneration. To characterize the molecular mechanism of SARM1 activation, we sought to determine the crystal structure of the SARM1 ARM domain. Here we describe the recombinant production and crystallization of dSARM1^ARM^, as well as unconventional process used for structure determination. Crystals were obtained in the presence of NMN, a precursor of NAD^+^ and a potential activator of SARM1, only after *in situ* proteolysis of the N-terminal 63 residues. After molecular replacement attempts failed, we determined the crystal structure of dSARM1^ARM^ at 1.65 Å resolution using the MIRAS phasing technique with the program autoSHARP, combining data from the native, SeMet-labelled, and Br-soaked crystals. The structure will further our understanding of the regulation of SARM1.

## 1. Introduction

The protein SARM1 (sterile alpha and Toll/interleukin-1 receptor motif-containing 1) is a central executioner of injury-induced axon degeneration (Wallerian degeneration). Loss of SARM1 protects axons from degeneration for weeks after injury induced by axotomy or vincristine (Osterloh et al., 2012, Gerdts et al., 2013). In healthy axons, SARM1 is held in the inactive state by the N-terminal ARM (armadillo repeat motif) domain. Upon injury, this auto-inhibition is relieved, permitting the C-terminal TIR (Toll/interleukin-1 receptor) domains to hydrolyze NAD^+^ (nicotinamide adenine dinucleotide) into nicotinamide, and either ADPR (ADP-ribose) or cADPR (cyclic ADPR) (Essuman et al., 2017, Horsefield et al., 2019). These changes in turn trigger an influx of Ca^2+^ to the cells, a corresponding loss of ATP and eventually axon degeneration (Loreto et al., 2015, Horsefield et al., 2019). Despite the important role in this process, the mechanism of SARM1 activation is poorly understood. Recently, it has been suggested that the accumulation of the NAD^+^ precursor NMN (nicotinamide mononucleotide) is a trigger of SARM1 activation, resulting in subsequent axon degeneration (Di Stefano et al., 2015). Therefore, we hypothesized that NMN may interact with the ARM domain and reverse the ARM domain-mediated auto-inhibition.

To determine crystal structures, the phase components of the structure factors, which are lost during diffraction data collection, need to be recovered. In the case where similar protein structures are available, one can use molecular replacement (Rossmann, 1990). Alternatively, one can locate the positions of heavy atoms (HAs) that are either incorporated into the crystals or already present within the macromolecule, through techniques such as MIR (multiple isomorphous replacement), usually with the inclusion of the anomalous signal, and SAD (single-wavelength anomalous dispersion) (Vijayan and Ramaseshan, 2001).

We sought to determine the crystal structure of the NMN-bound *Drosophila* SARM1 ARM domain (dSARM1^ARM^), which would greatly enhance our understanding of the mechanism of NMN-induced relief of ARM domain-mediated auto-inhibition in SARM1. We crystallized dSARM1^ARM^ in the NMN-bound state. Although the crystals diffracted X-rays to high resolution (1.65 Å), attempts to determine the phases using molecular replacement technique were not successful. We then attempted SAD phasing using the anomalous signals from bromide (Br) or selenium (Se) atoms, which were separately incorporated into the crystals. However, the anomalous signal present in either the Br-SAD or Se-SAD dataset was weak and initial phases could not be successfully estimated. The phase problem was eventually solved by employing the MIRAS (multiple isomorphous replacement with anomalous scattering) method with the program autoSHARP (Vonrhein et al., 2007), combining the data from the native, the SeMet (selenomethionine)-labelled, and the Br-soaked crystals. Here, we report the protein production, crystallization, and structure determination of the dSARM1^ARM^, and present our experience as a case study of modern MIRAS phasing.

## 2. Materials and methods

### 2.1. Protein production

The cDNA encoding dSARM1^ARM^ (residues 307-678; UniProtKB: Q6IDD9) was codon-optimized for expression in *Escherichia coli* (*E. coli*), and cloned into the the pMCSG7 expression vector at the *SspI* site, using ligation-independent cloning technique (forward primer: 5’-TACTTCCAATCCAATGCGAATGGACAGATGTTGAAGCTTGCGGATTTGAAATTAGACG-3’; reverse primer: 5’-TTATCCACTTCCAATGTTACGTTTCCCCAATTAAGCGCAGCGCTTGGG-3’) (Aslanidis and de Jong, 1990). The plasmid was transformed into *E. coli* BL21 (DE3) (for native protein) or B834 (DE3) cells (for SeMet-labelled protein) by heat shock. Cells were grown on LB (lysogeny broth) agar plates containing 100 μg/mL ampicillin at 37 °C overnight. Colonies were inoculated into 10 mL of LB media containing 100 μg/mL ampicillin, and incubated at 37 °C, 225 rpm overnight. To produce native protein, 1 mL of the LB overnight culture of transformed *E. coli* BL21 (DE3) cells was inoculated into 1000 mL of auto-induction media (Studier, 2005) containing 100 μg/mL ampicillin and incubated at 37 °C, 225 rpm until OD_600_ reached 0.8-1.0. The temperature was then decreased to 20 °C for overnight protein expression. To produce SeMet-labelled protein, 1 mL of the LB overnight culture of transformed *E. coli* B834 (DE3) cells was inoculated into l000 mL of M9 minimal media containing 1× M9 salt (33.7 mM Na_2_HPO_4_, 22 mM KH_2_PO_4_, 8.55 mM NaCl and 9.35 mM NH_4_Cl), 1× trace elements solution (0.13 mM EDTA, 0.03 mM FeCl_3_, 6.2 μM ZnCl_2_, 0.76 μM CuCl_2_, 0.42 μM CoCl_2_, 1.62 μM H_3_BO_3_ and 0.08 μM MnCl_2_), 0.4% (v/v) glucose, 1 mM MgSO_4_, 0.3 mM CaCl_2_, 1× BME vitamin solution (Sigma-Aldrich), and 100 μg/mL ampicillin. The bacteria were grown at 37 °C, 225 rpm until OD_600_ reached 0.8-1.0. The temperature was then decreased to 20 °C for a 30 min incubation. One mL of 50 mg/mL SeMet (Sigma-Aldrich) was added to 1000 mL of culture. The expression was induced by adding IPTG (isopropyl β-D-1-thiogalactopyranoside) to a final concentration of 1 mM and the cells were incubated overnight at 20 °C, 225 rpm. *E. coli* BL21 (DE3) or B834 (DE3) cells were harvested by centrifugation at 6000 × *g* for 20 min at 4 °C, and were treated identically in subsequent purification steps.

The harvested cells were resuspended in lysis buffer (50 mM HEPES (pH 8.0), 500 mM NaCl, 30 mM imidazole and 1 mM DTT). Phenymethylsulfonyl fluoride was added to the cell suspension at a final concentration of 1 mM. Cells were subsequently lysed by sonication. The lysed cells were centrifuged at 15300 × *g* for 40 min at 4 °C. The supernatant was loaded onto a 5 mL HisTrap HP column (Cytiva) equilibrated with lysis buffer. The bound target protein was washed with 100 mL of lysis buffer and eluted with elution buffer (50 mM HEPES (pH 8.0), 500 mM NaCl, 300 mM imidazole, and 1 mM DTT) on an AKTA Purifier (Cytiva). Fractions containing dSARM1^ARM^ were combined and incubated with TEV (tobacco etch virus) protease (protein: TEV protease = 20:1), in the SnakeSkin Dialysis Tubing, 3.5 kDa MWCO (Thermo Fisher Scientific) and dialyzed against buffer containing 20 mM HEPES (pH 8.0), 300 mM NaCl and 1 mM DTT at 4 °C overnight. The His_6_-tag-removed protein was re-loaded onto a 5 mL HisTrap HP column to remove uncleaved fusion protein and free His_6_ tag. The flow-through was then collected and concentrated to 10 mL, and injected onto a Superdex 75 HiLoad 26/600 column (Cytiva) equilibrated with gel filtration buffer (10 mM HEPES (pH 8.0), 150 mM NaCl, and 1mM DTT). The peak fractions containing pure dSARM1^ARM^ were pooled, concentrated using a 30 kDa MWCO Amicon Ultra Centrifugal filter (Millipore), flash-frozen, and stored at −80 °C.

### 2.2. Crystallization

Prior to crystallization, dSARM1^ARM^ protein (17 mg/mL) was incubated with NMN in a 1:10 protein-to-compound molar ratio at 4 °C overnight. Sparse matrix protein crystallization screening was performed using the commercially available screens Index (Hampton Research), Combined Synergy (Hampton Research), PEG/Ion (Hampton Research), PEGRx (Hampton Research), JCSG+ (Molecular Dimension), PACT Premier™ (Molecular Dimension), ProPlex™ (Molecular Dimension), and ShotGun (Molecular Dimension). Crystallization trials were set up using a Mosquito liquid handling robot (TTP LabTech) in a 96-well hanging-drop plate format with 100 nL protein and 100 nL reservoir solution per drop, equilibrated against 75 μL of reservoir solution at 20 °C. To scale up the drop sizes, hanging drops consisting of 2 μL of NMN-bound protein and 2 μL of commercial reservoir solution were equilibrated against 500 μL of home-made reservoir solution containing 0.1 M SPG buffer (succinic acid, sodium dihydrogen phosphate, and glycine in the molar ratios 2:7:7; pH 8.0) and 25% (w/v) PEG1500 at 20 °C. Crystals were observed in the drops after 3-5 days. SDS-PAGE and mass spectrometry analyses of these crystals were performed to ascertain the identity of the crystallized protein.

### 2.3. Diffraction data collection and processing

Prior to flash-cooling in liquid nitrogen, crystals of NMN-bound native dSARM1^ARM^ and SeMet-labelled dSARM1^ARM^ were cryo-protected in a cryoprotectant solution containing 0.1 M SPG buffer (pH 8.0), 25% (w/v) PEG 1500, and 25% (v/v) PEG 400. Crystals derivatized with Br were prepared by soaking the native crystals in the mother liquor containing 0.5 M sodium bromide and 25% PEG 400 (v/v) for 2 min prior to flash-cooling. All datasets were collected at the Australian Synchrotron on the MX2 beamline (Aragão et al., 2018). The native dataset was collected at a wavelength of 0.95372 Å, and the SeMet-labelled dataset was collected at the theoretical Se absorption edge with a wavelength of 0.97857 Å., and the Br-soaked dataset was collected at the theoretical Br absorption edge with a wavelength of 0.91976 Å. Diffraction data of NMN-bound dSARM1^ARM^ crystals (native, SeMet-labelled and Br-soaked data) were processed and analyzed with the program autoPROC (Vonrhein et al., 2011). Initial phases were calculated using MIRAS technique, with the program autoSHARP (Vonrhein et al., 2007). The structure was refined using iterations of phenix.refine (Afonine et al., 2012) and manual model building in Coot (Emsley and Cowtan, 2004).

## 3. Results and discussion

The boundaries of the expression constructs for dSARM1^ARM^ were determined based on sequence alignments among human, mouse, *Caenorhabditis elegans*, and *Drosophila* SARM1, taking into consideration secondary structure predictions. Small-scale expression tests were performed to identify constructs producing soluble target protein. Using the dSARM1^ARM307-678^ construct, we successfully expressed soluble dSARM1^ARM^ protein with a final yield of 10 mg/liter of bacterial culture.

During sparse matrix screening, we observed (after 5 days) a few crystals grown in 0.1 M SPG buffer (pH 8.0) and 25% (w/v) PEG1500 (PACT premier™ condition A5) at 20 °C. The crystals were chunky but irregular in shape (Figure 1). However, despite our best efforts, we were not able to reproduce NMN-bound dSARM1^ARM^ crystals using home-made crystallization solutions.

**Figure 1:**
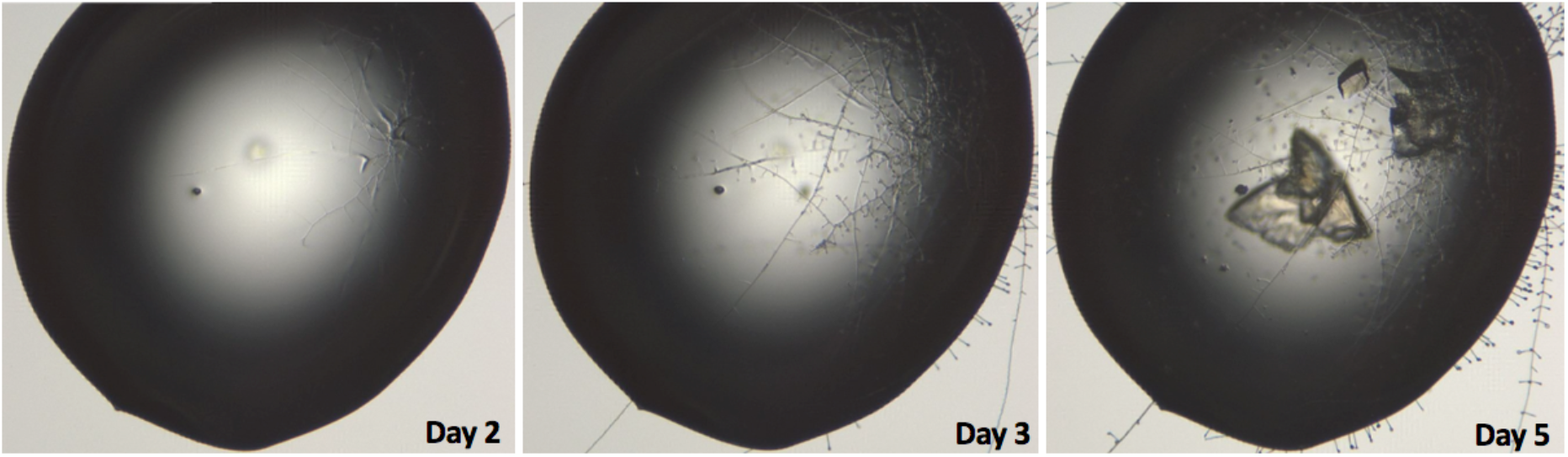
Crystal growth of NMN-bound dSARM1^ARM^. Crystals were observed after 5 days in 0.1 M SPG buffer (pH 8.0) and 25% (w/v) PEG 1500 in the presence of fungal cells.

Importantly, we observed what appeared to be fungal cells, possibly *Penicillium*, grew in the crystallization drop prior to the growth of dSARM1^ARM^ crystals (Figure 1). SDS-PAGE analysis of the NMN-bound dSARM1^ARM^ crystals, followed by mass spectrometry, indicated that the crystallized protein corresponded to residues 370-678 of SARM1 (i.e. lacking the N-terminal 63 amino-acids). This led us to reason that partial proteolysis, mediated by proteases secreted by the fungal cells, was required for NMN-bound SARM1^ARM^ crystallization. We therefore produced dSARM1^ARM370-678^, in the hope of solving the crystal reproducibility issues, however, this construct failed to yield soluble protein. Crystals only grew when the original crystallization solution containing the fungal cells was added to the home-made crystallization solution. For these reasons, we had a limited number of crystals to optimize our diffraction experiments.

A native dataset from NMN-bound dSARM1^ARM^ crystals was collected at the Australian Synchrotron on the MX2 beamline. The crystals diffracted to > 1.65 Å resolution (Figure 2, Table 1). The crystal had the symmetry of the space group *P*1 and likely contained two dSARM1^ARM^ molecules in the ASU (asymmetric unit), with a Matthews coefficient of 2.07 Å^3^/Dalton and a solvent content of 40 %.

**Figure 2:**
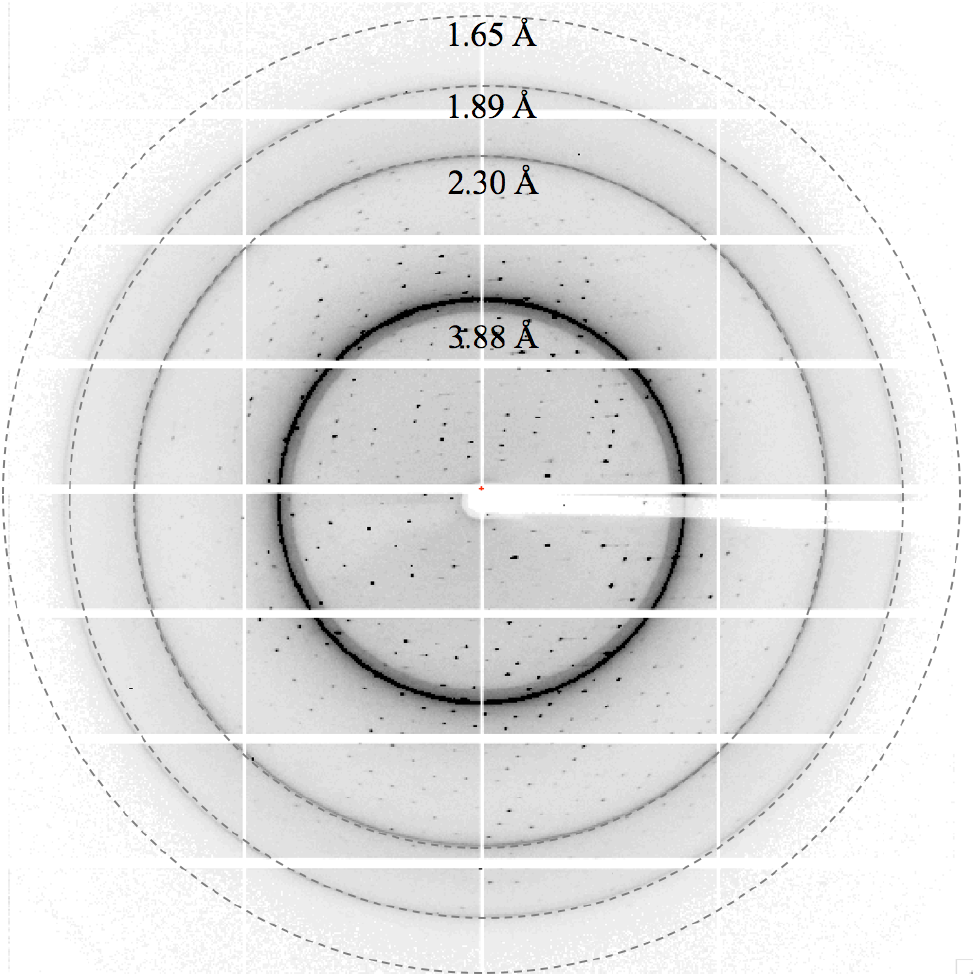
A representative diffraction image of NMN-bound dSARM1^ARM^ crystals. The crystal diffracted X-rays to >1.65 Å resolution.

**Table 1:** Data processing statistics from XDS and Aimless.

| Data collection |  |  |  |
| --- | --- | --- | --- |
|  | Native | Br-soaked | SeMet-labelled |
| Space group | P1 | P1 | P1 |
| a, b, c (Å) | 38.95, 50.82, 76.12 | 39.12, 51.15, 75.83 | 38.86, 50.28, 75.18 |
| $\alpha, \beta, \gamma$ (°) | 103.52, 101.89, 95.23 | 103.37, 101.51, 96.67 | 104.88, 101.37, 94.89 |
| Resolution (Å) | 48.9-1.74 (1.78-1.74) | 46.59-2.01 (2.06-2.01) | 48.1-4.18 (4.67-4.18) |
| R<sub>merge</sub> | 0.058 (0.472) | 0.067 (0.621) | 0.036 (0.043) |
| R<sub>meas</sub> | 0.068 (0.554) | 0.077 (0.719) | 0.051 (0.060) |
| R<sub>pim</sub> | 0.035 (0.288) | 0.038 (0.357) | 0.036 (0.043) |
| Mean I/σ (I) | 17.6 (3.4) | 17.8 (3.2) | 21.7 (19.9) |
| CC<sub>1/2</sub> | 0.999 (0.931) | 0.999 (0.830) | 0.98 (1.13) |
| Total number of observations | 3386,169 (19,534) | 284,357 (19,417) | 13,961 (3,827) |
| Total number unique | 54,045 (2,793) | 35,879 (2,561) | 3,835 (1,074) |
| Completeness (%) | 96.3 (89.2) | 97.9 (93.5) | 97.9 (96.8) |
| Multiplicity | 7.1 (7.0) | 7.9 (7.6) | 3.6 (3.6) |
| Anomalous completeness (%) | 95.4 (85.5) | 95.3 (88.5) | 92.6 (89.4) |
| Anomalous multiplicity | 3.5 (3.6) | 3.9 (3.9) | 1.8 (1.9) |
| DelAnom correlation between half-sets | -0.091 (0.076) | 0.398 (0.053) | 0.297 (0.183) |
| Mid-slope of anomalous normal probability | 0.956 | 1.173 | 1.392 |

1. The statistics are based on the calculations from Aimless. 2. The numbers in parentheses represent the highest resolution shell. 3. $R_{\text{merge}} = \sum_{hkl} \sum_j |I_{hkl,j} - \langle I_{hkl} \rangle| / (\sum_{hkl} \sum_j I_{hkl,j})$ ; $R_{\text{meas}} = \sum_{hkl} [N/(N-1)]^{1/2} \sum_j |I_{hkl,j} - \langle I_{hkl} \rangle| / (\sum_{hkl} \sum_j I_{hkl,j})$ ; $R_{\text{pim}} = \sum_{hkl} [1/(N-1)]^{1/2} \sum_j |I_{hkl,j} - \langle I_{hkl} \rangle| / (\sum_{hkl} \sum_j I_{hkl,j})$

ARM domains are protein interaction domains found in many proteins displaying diverse cellular roles from gene expression to cytoskeleton regulation (Coates, 2003). They typically consist of tandem repeats of armadillo motifs. One armadillo motif contains ~42 residues, which fold into three alpha helices (H1, H2 and H3). Stacking of these motifs forms a right-handed superhelix with an elongated concave surface, characterized by parallel H3 helices arranged in a ladder fashion (Coates, 2003). Initially, we attempted to solve the structure of dSARM1^ARM^ by molecular replacement. We used the available ARM structures, such as importin-α (PDB: 1IAL) (Kobe, 1999) and Vac8p (PDB: 5XJG) (Jeong et al., 2017), with various modifications, as search models, within the program Phaser (McCoy et al., 2007). We attempted automated molecular replacement with the programs MrBUMP (Keegan and Winn, 2008) and Balbes (Long et al., 2008), as well as *ab initio* macromolecular phasing with the tool ARCIMBOLDO (Rodríguez et al., 2009). However, we did not obtain any clear solutions using these approaches.

We alternatively sought to solve the phase problem by SAD phasing. To this end, we incorporated Br (through soaking), and separately, SeMet (during expression), into the protein crystals and collected anomalous datasets to 2.0 Å and 4.2 Å resolution, respectively, using X-ray wavelengths close to the absorption edges of the respective HAs (Table 1). Unfortunately, the crystals were prone to radiation damage, as demonstrated by the reduction in diffraction quality and a sharply decreasing Wilson B factor. Due to the low *P*1 symmetry, datasets with high multiplicity and therefore accurately determined anomalous differences, which are often required for successful SAD phasing, could not be acquired. Also, as we only had access to a limited number of crystals, merging multiple low-multiplicity datasets was not a viable option. The Br dataset had an anomalous multiplicity of ~3.9, whereas the SeMet dataset had an even lower anomalous multiplicity of merely 1.8. Data processing using XDS (Kabsch, 2010) and Aimless (Evans and Murshudov, 2013) indicated that the values of the mid-slope of anomalous normal probabilities of the Br and SeMet datasets were 1.17 and 1.39, respectively, suggesting that detectable but weak anomalous signal was present in these two datasets. Using a CC_ano_ cutoff of 0.15, the detectable anomalous signals were up to 2.7 Å and 4.2 Å for the Br and SeMet dataset, respectively (Figure 4). However, subsequent searches for HAs invariably failed.

Fortunately, the Br-soaked and SeMet-labelled crystals had similar unit cells dimensions to the native protein crystals (Table 1), indicating that they were probably all quite isomorphous to each other, thus making MIRAS or SIRAS (single isomorphous replacement with anomalous scattering) phasing a possibility. Therefore, we employed the software package autoPROC for data reprocessing (Vonrhein et al., 2011), which uses XDS for data processing (Kabsch, 2010), POINTLESS for space-group determination (Evans, 2006), AIMLESS for scaling (Evans and Murshudov, 2013) and STARANISO for analysis of diffraction anisotropy (Vonrhein et al., 2018), plus multiple additional tools for handling diffraction data processing (Table 2, Figure 5). For MIRAS phasing and initial model building, we used autoSHARP (Vonrhein et al., 2007), which uses SHELXC/D for substructure determination (Schneider and Sheldrick, 2002), SHARP for HA refinement, phasing and completion (de La Fortelle and Bricogne, 1997), SOLOMON for density modification (Abrahams and Leslie, 1996), and BUCCANEER for automatic model building (Cowtan, 2006). Careful processing of the diffraction images is important, as outliers are damaging to the success rate of the HA search. Using autoPROC, we reprocessed the raw data of the native, Br, and the SeMet datasets (Table 2). To ensure that no anomalous differences - especially at low resolution - were affected and could hinder successful substructure detection, we inspected the diffraction images visually to define accurate beam-stop masks for processing. We also identified and removed damaged pixels in the SeMet dataset using the tools provided by autoPROC, which proved to be critical for successfully processing the SeMet dataset.

**Table 2:** Data processing statistics from autoPROC and refinement statistics from Phenix.

| Data collection |  |  |  |
| --- | --- | --- | --- |
|  | Native | Br-soaked | SeMet-labelled |
| Space group | P1 | P1 | P1 |
| a, b, c (Å) | 38.92, 50.79, 76.05 | 39.07, 51.08, 75.73 | 38.89, 50.31, 75.22 |
| $\alpha, \beta, \gamma$ (°) | 103.52, 101.90, 95.26 | 103.38, 101.52, 96.66 | 104.86, 101.36, 94.91 |
| Resolution (Å) | 71.9-1.46 (1.60-1.46) | 49.0-1.68 (1.82-1.68) | 48.1-1.89 (2.11-1.89) |
| R<sub>merge</sub> | 0.05 (0.95) | 0.08 (1.77) | 0.07 (0.76) |
| R<sub>meas</sub> | 0.06 (1.15) | 0.09 (2.05) | 0.09 (1.08) |
| R<sub>pim</sub> | 0.03 (0.63) | 0.04 (1.01) | 0.07 (0.76) |
| Mean I/σ (I) | 15.9 (1.6) | 13.9 (1.0) | 9.2 (1.2) |
| CC<sub>1/2</sub> | 1.00 (0.63) | 1.00 (0.4) | 1.00 (0.47) |
| Total reflections | 456,719 (20,265) | 393,818 (19,421) | 86,502 (4,041) |
| Unique reflections | 64,028 (3,201) | 48,934 (2,447) | 24,291 (1,216) |
| Completeness (spherical) | 67.2 (14.2) | 78.2 (19.1) | 56.6 (10.0) |
| Completeness (ellipsoidal) | 83.5 (40.3) | 90.0 (44.7) | 85.3 (38.8) |
| Multiplicity | 7.1 (6.3) | 8.0 (7.9) | 3.6 (3.3) |
| Anomalous completeness (spherical) | 66.5 (14.0) | 76.4 (18.5) | 55.8 (9.7) |
| Anomalous completeness (ellipsoidal) | 82.7 (39.8) | 87.9 (43.4) | 84.1 (38.0) |
| Anomalous multiplicity | 3.6 (3.2) | 4.1 (4.1) | 1.8 (1.7) |
| Refinement |  |  |  |
| Resolution (Å) | 37.67-1.65 (native dataset) |  |  |
| R<sub>work</sub> | 0.21 |  |  |
| R<sub>free</sub> | 0.23 |  |  |
| RMS bonds (Å) | 0.002 |  |  |
| RMS angles (°) | 0.47 |  |  |
| Ramachandran favored (%) | 98.51 |  |  |
| Ramachandran outliers (%) | 0.33 |  |  |
| Rotamer outliers (%) | 0.19 |  |  |
| Clashscore | 1.64 |  |  |
| Average B-factor (Å<sup>2</sup>) | 27.77 |  |  |
| C-beta outliers | 0 |  |  |

1. The statistics are based on the calculations from autoPROC and MolProbity. 2. The numbers in parentheses represent the highest resolution shell. 3. $R_{\text{merge}} = \sum_{\text{hkl}} \sum_j |I_{\text{hkl},j} - \langle I_{\text{hkl}} \rangle| / (\sum_{\text{hkl}} \sum_j I_{\text{hkl},j})$ ; $R_{\text{meas}} = \sum_{\text{hkl}} [N/(N-1)]^{1/2} \sum_j |I_{\text{hkl},j} - \langle I_{\text{hkl}} \rangle| / (\sum_{\text{hkl}} \sum_j I_{\text{hkl},j})$ ; $R_{\text{pim}} = \sum_{\text{hkl}} [1/(N-1)]^{1/2} \sum_j |I_{\text{hkl},j} - \langle I_{\text{hkl}} \rangle| / (\sum_{\text{hkl}} \sum_j I_{\text{hkl},j})$ 4. $R_{\text{work}} = \sum_{\text{hkl}} |F_{\text{obs},\text{hkl}} - F_{\text{calc},\text{hkl}}| / \sum |F_{\text{obs},\text{hkl}}|$ ; $R_{\text{free}}$ is equivalent to $R_{\text{work}}$ , with 5% of data excluded from refinement process. $|F_{\text{obs},\text{hkl}}|$ and $|F_{\text{calc},\text{hkl}}|$ represent the observed and calculated structure factor amplitudes.

Phase determination was performed using the program autoSHARP (Vonrhein et al., 2007), with a resolution cutoff of 25-2.5 Å. The high-resolution cutoff seems slightly counterintuitive, because it is (1) much higher than the HA signal in the different datasets and (2) lower than the overall diffraction limit of the available datasets. However, in cases where the difference between those two resolution limits (HA signal and overall) is rather large, it is often beneficial to restrict the data to 2.5-3 Å at the early stages of the structure solution process. At this stage, one is mainly interested in achieving a successful substructure determination, a clear indication of the correct enantiomorph (during the density modification step), and hopefully some meaningful secondary structure elements resulting from the automatic model building step, all this can easily be achieved by using 2.5-3 Å data. If even higher resolution data is used at this stage, the initial low-resolution (and expected to be poor) phase information might be inadequate in helping density modification bridging that large resolution range to the full limit of the available data. During HA detection - where several combinations of MIRAS (native, Br and SeMet datasets), SIRAS (native plus Br or native plus SeMet datasets) and SAD (Br or SeMet dataset alone) were tried, the solution with the highest correlation coefficient between the observed and calculated E values (normalized substructure amplitudes) corresponded to SIRAS of SeMet (CC(E) = 0.188 with 4 SeMet sites out of the expected 10 sites in the ASU). Using this solution as a starting point, SHARP within autoSHARP further detected 10 Br sites (overall phasing power of 0.216 for isomorphous differences and 0.483 for anomalous differences, with the phasing power dropping below one at 24.42 Å and 5.48 Å, respectively) and 10 SeMet sites (overall phasing power of 0.136 for isomorphous differences and 0.299 for anomalous differences, with the phasing power dropping below one at 24.42 Å and 4.92 Å, respectively). These values were consistent with the analysis of the anomalous signal from the data processing stages and confirmed that the HA signal (and initial maps computed with those phases) would present a rather low resolution starting point for subsequent steps. A final set of phases was calculated in both hands and the most likely enantiomorph was determined by performing a single cycle of solvent flipping in SOLOMON as part of autoSHARP, suggesting that the correct phases were those from the inverted hand, based on its slightly higher score (a combination of the correlation coefficient between observed E^2^ values and E^2^ values of the modified map and the contrast in the assigned protein and solvent regions) of 0.1234 (2 molecules/ASU), against 0.1078 from the original hand. After multiple cycles of density modification to optimize the solvent content, the best density-modified map with a score of 1.8697 was finally handed over to BUCCANEER, which managed to build a total of 614 residues in two chains (out of the expected 618 residues for a dimer in the ASU). This was the final result of a fully automatic autoSHARP run (starting with the datasets, the sequence, information about the scattering properties of the different datasets and some indication of the expected number of Br sites) and provided the starting point for subsequent steps. Further manual model building in Coot (Emsley and Cowtan, 2004) and refinement using phenix.refine (Afonine et al., 2012) improved the quality of phases (Figure 3). The final structure was determined with R_work_ and R_free_ values of 0.21 and 0.23, respectively (Table 2).

**Figure 3:**
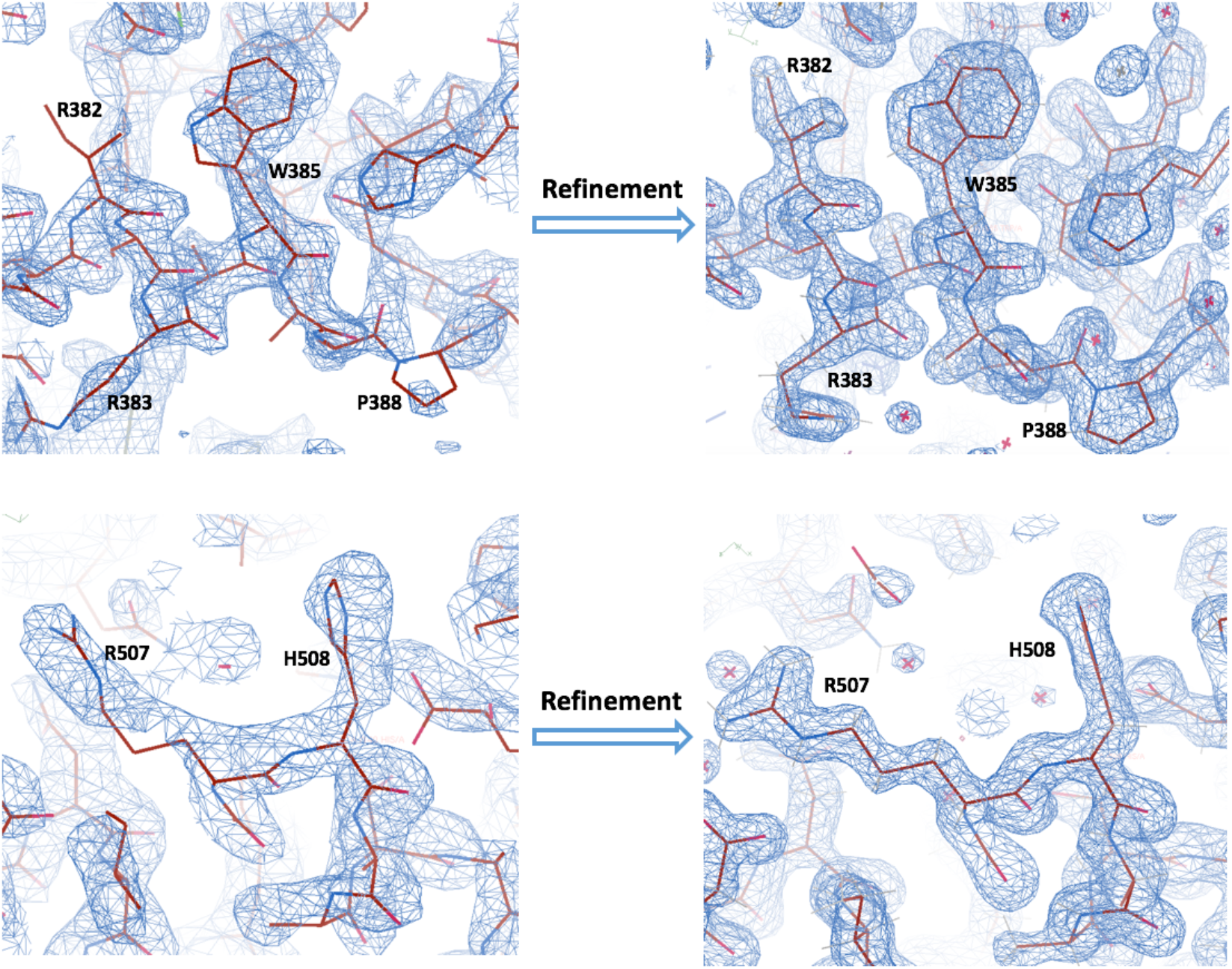
Representative electron density maps, contoured at 1.5 σ, before and after the refinement. Panels on the left show the initial electron density map (2F_O_-F_C_), calculated using the MIRAS-based phases from autoSHARP after density modification (SOLOMON) and model building (BUCCANEER) at a resolution of 2.5 Å. Panels on the right show the electron density map in the corresponding region after refinement with phenix.refine at 1.65 Å resolution.

**Figure 4:**
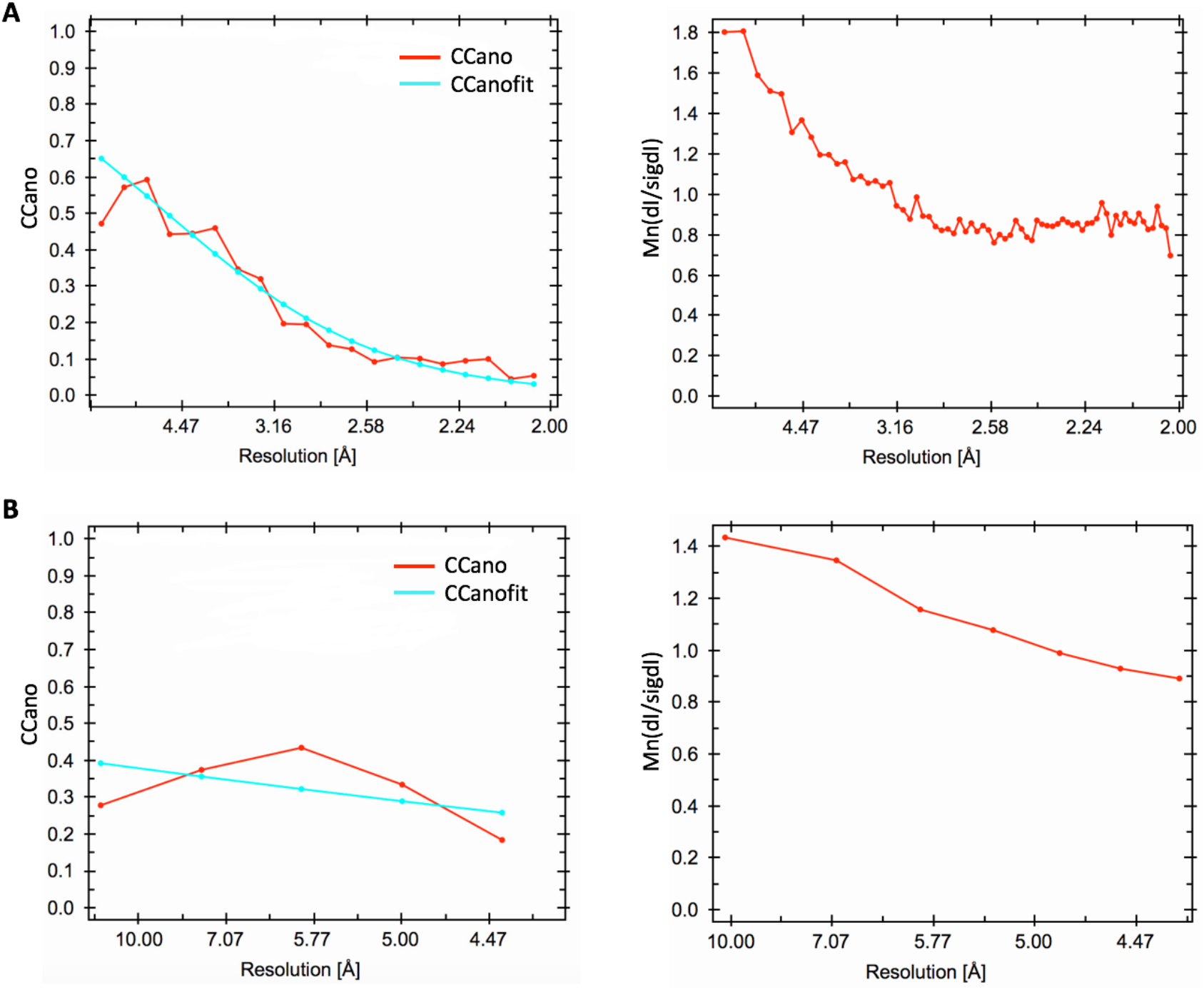
Analysis of the anomalous signals using XDS. A. CCano (left) and Mn(dI/sigdI) (right) as a function of resolution for the Br dataset. B. CCano (left) and Mn(dI/sigdI) (right) as a function of resolution for the SeMet dataset. The figures were automatically generated during data processing.

**Figure 5:**
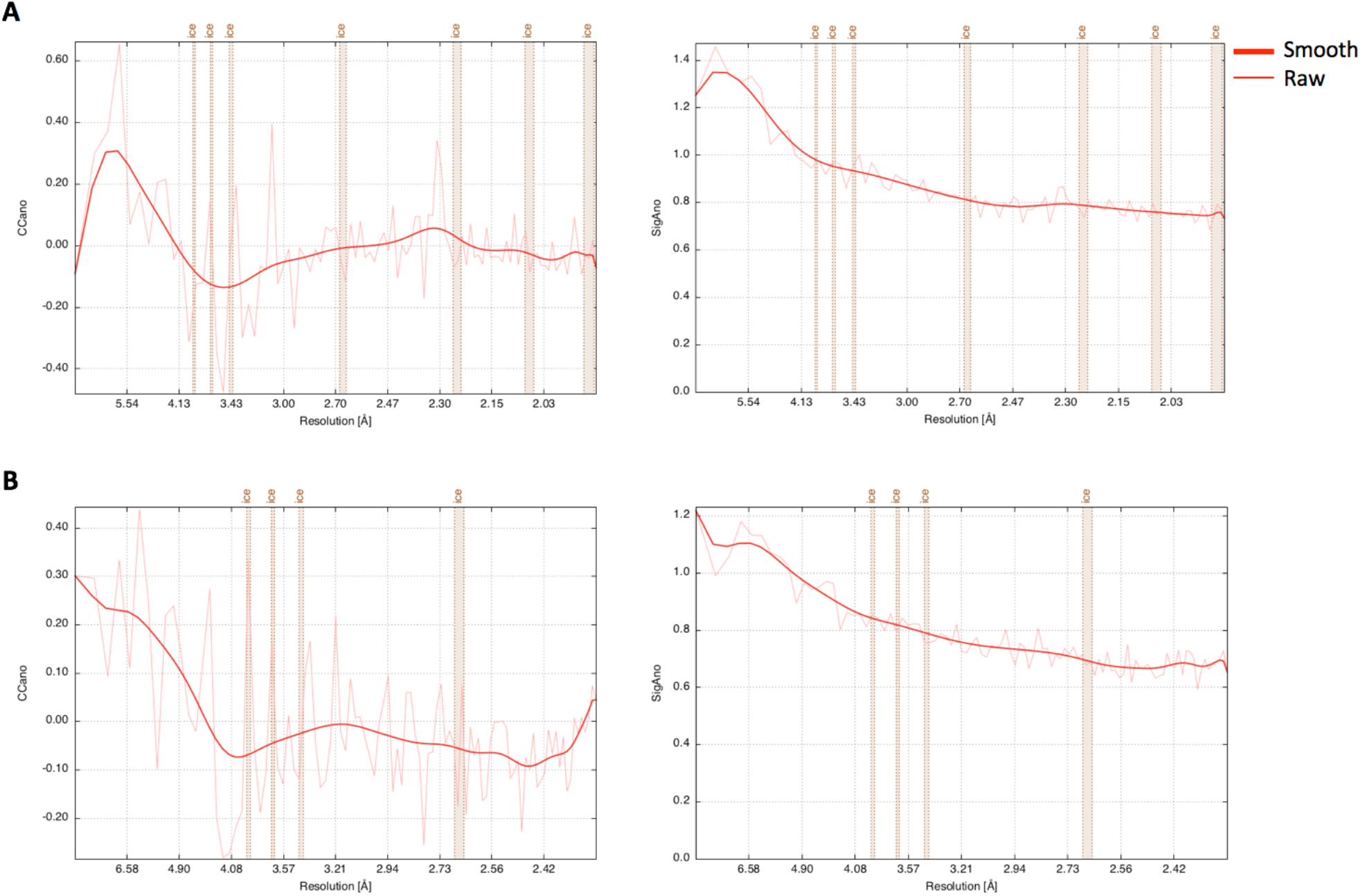
Analysis of the anomalous signals using autoPROC. A. CCano (left) and SigAno (right) as a function of resolution for the Br dataset. B. CCano (left) and SigAno (right) as a function of resolution for the SeMet dataset. The figures were automatically generated during data processing.

## 4. Summary

In conclusion, we describe the crystal structure determination of dSARM1^ARM^, as an illustrative example of how a number of technical difficulties in the process can be overcome. A number of non-conventional steps were employed here. First, *in situ* proteolysis allowed diffraction-quality crystals to be obtained in the first place. Second, after attempts to solve the phase problem by molecular replacement failed, HAs were introduced by Br soaking and SeMet incorporation. However, neither of these SAD datasets provided sufficient anomalous signal to obtain interpretable electron density maps. The low symmetry of the crystal system provided little opportunity to collect high-multiplicity multi-wavelength datasets around the Se and Br edges, which might have helped substructure detection within each Br or SeMet dataset. However, after careful data processing (manual beam-stop masking, exclusion of damaged pixels that are not yet in the detector mask and handling of ice-rings) using autoPROC and combining the resulting improved data from the native, Br-soaked and SeMet-labelled crystals, the MIRAS approach as implemented in autoSHARP led to interpretable electron density maps and a clear initial starting model. This structure will help us understand the molecular mechanisms of regulation of SARM1, a protein with a central role in neurodegenerative disease. The biological implications of the structure will be discussed elsewhere.

## 5. Acknowledgements

We acknowledge the use of the University of Queensland Remote Operation Crystallization and X-ray (UQROCX) Facility at the Centre for Microscopy and Microanalysis and the support from staff, Dr. Gordon King and Karl Byriel. We acknowledge the use of the Australian Synchrotron MX beamlines, part of ANSTO, including the Australian Cancer Research Foundation detector, and the support of staff. We also thank the CCP4/Shanghai 2019 workshop, their organisers, including CCP4, Dr. Ruslan Sanishvili, National Facility for Protein Science in Shanghai, Shanghai Tech University iHuman Institute, and Shanghai Synchrotron Radiation Facility, and Prof. Gérard Bricogne from Global Phasing Ltd for the help with structural determination. We are grateful to Dr Gayle Petersen for diligent proofreading, and discussion of the manuscript. This work was supported by grants from the Australian National Health and Medical Research Council (NHMRC; 1107804 and 1160570 to BK and TV, 1071659 to BK and 1108859 to TV) and the Australian Research Council (ARC; Laureate Fellowship FL180100109 to BK). TV received ARC Discovery Early Career Researcher Award (DECRA) funding (DE170100783).

## References

Abrahams, J. P. & Leslie, A. G. (1996). Acta Cryst. D52, 30–42.

Afonine, P. V., Grosse-Kunstleve, R. W., Echols, N., Headd, J. J., Moriarty, N. W., Mustyakimov, M., Terwilliger, T. C., Urzhumtsev, A., Zwart, P. H. & Adams, P. D. (2012). Acta Cryst. D68, 352–367.

Aragão, D., Aishima, J., Cherukuvada, H., Clarken, R., Clift, M., Cowieson, N. P., Ericsson, D. J., Gee, C. L., Macedo, S., Mudie, N., Panjikar, S., Price, J. R., Riboldi-Tunnicliffe, A., Rostan, R., Williamson, R. & Caradoc-Davies, T. T. (2018). J Synchrotron Radiat. 25, 885–891.

Aslanidis, C. & de Jong, P. J. (1990). Nucleic Acids Res. 18, 6069–6074.

Coates, J. C. (2003). Trends Cell Biol. 13, 463–471.

Cowtan, K. (2006). Acta Cryst. D62, 1002–1011.

de La Fortelle, E. & Bricogne, G. (1997). Maximum-likelihood heavy-atom parameter refinement for multiple isomorphous replacement and multiwavelength anomalous diffraction methods. pp. 472–494. Academic Press.

Di Stefano, M., Nascimento-Ferreira, I., Orsomando, G., Mori, V., Gilley, J., Brown, R., Janeckova, L., Vargas, M. E., Worrell, L. A., Loreto, A., Tickle, J., Patrick, J., Webster, J. R. M., Marangoni, M., Carpi, F. M., Pucciarelli, S., Rossi, F., Meng, W., Sagasti, A., Ribchester, R. R., Magni, G., Coleman, M. P. & Conforti, L. (2015). Cell Death Differ. 22, 731–742.

Emsley, P. & Cowtan, K. D. (2004). Acta Cryst. D60, 2126–2132.

Eschenfeldt, W. H., Lucy, S., Millard, C. S., Joachimiak, A. & Mark, I. D. (2009). Methods Mol Biol. 498, 105–115.

Essuman, K., Summers, D. W., Sasaki, Y., Mao, X., Diantonio, A. & Milbrandt, J. (2017). Neuron. 93, 1334–1343.

Evans, P. (2006). Acta Cryst. D62, 72–82.

Evans, P. R. & Murshudov, G. N. (2013). How good are my data and what is the resolution? Acta Cryst. D69, 1204–1214.

Gerdts, J., Summers, D. W., Sasaki, Y., Diantonio, A. & Milbrandt, J. (2013). J Neurosci. 33, 13569.

Horsefield, S., Burdett, H., Zhang, X., Manik, M. K., Shi, Y., Chen, J., Qi, T., Gilley, J., Lai, J.S., Rank, M. X., Casey, L. W., Gu, W., Ericsson, D. J., Foley, G., Hughes, R. O., Bosanac, T., Von Itzstein, M., Rathjen, J. P., Nanson, J. D., Boden, M., Dry, I. B., Williams, S. J., Staskawicz, B. J., Coleman, M. P., Ve, T., Dodds, P. N. & Kobe, B. (2019). Science. 365, 793–799.

Jeong, H., Park, J., Kim, H. I., Lee, M., Ko, Y. J., Lee, S., Jun, Y. & Lee, C. (2017). Proc Nati Acad Sci. 114, E4539–E4548.

Kobe, B. (1999). Nat Struct Biol. 6, 388–397.

Loreto, A., Di Stefano, M., Gering, M. & Conforti, L. (2015). Cell Rep. 13, 2539–2552.

Long, F., Vagin, A. A., Young, P. & Murshudov, G. N. (2008). Acta Cryst. D64, 125–132.

McCoy, A. J., Grosse-Kunstleve, R. W., Adams, P. D., Winn, M. D., Storoni, L. C. & Read, R. J. (2007). J Appl Cryst. 40, 658–674.

Osterloh, J. M., Yang, J., Rooney, T. M., Fox, A. N., Adalbert, R., Powell, E. H., Sheehan, A. E., Avery, M. A., Hackett, R., Logan, M. A., Macdonald, J. M., Ziegenfuss, J. S., Milde, S., Hou, Y. J., Nathan, C., Ding, A., Brown, R. H., JR., Conforti, L., Coleman, M., Tessier-Lavigne, M., Zuchner, S. & Freeman, M. R. (2012). Science. 337, 481–484.

Rodríguez, D. D., Grosse, C., Himmel, S., González, C., De Ilarduya, I. M., Becker, S., Sheldrick, G. M. & Usón, I. (2009). Nat Methods. 6, 651–653.

Rossmann, M. (1990). Acta Cryst. A46, 73–82.

Schneider, T. R. & Sheldrick, G. M. (2002). Acta Cryst. D58, 1772–1779.

Studier, F. W. (2005). Protein Expr Purif. 41, 207–234.

Vijayan, M. & Ramaseshan, S. (2001). Isomorphous replacement and anomalous scattering, edited by Shmueli, U, pp. 264–275. Dordrecht: Springer Netherlands.

Vonrhein, C., Blanc, E., Roversi, P. & Bricogne, G. (2007). Methods Mol Biol, 364, 215–230.

Vonrhein, C., Flensburg, C., Keller, P., Sharff, A., Smart, O., Paciorek, W., Womack, T. & Bricogne, G. (2011). Acta Cryst. D67, 293–302. a360.

